# Intranasal Delivery of HIV/SIV Antigens with NE/AS01B Adjuvants Enhances Cellular Immunity and Reduces Viral Loads in SHIV-Challenged Macaques

**DOI:** 10.64898/2026.02.04.703720

**Authors:** Michellie Thurman, Viswanathan Chokkavelu, Samuel D. Johnson, Omalla A Olwenyi, Gokul Raj Kathamuthu, Jianshi Yu, Samson Adeniji, Kai Ying Hong, Morgan Johnston, Deepanwita Bose, Kabita Pandey, Hongmei Gao, Xiaoying Shen, David Montefiori, Pamela T. Wong, James R Baker, Francois Villinger, Maureen Kane, Mohamed Abdel-Mohsen, Siddappa N. Byrareddy

## Abstract

The primary route of HIV transmission is across mucosal tissues; therefore, developing a protective mucosal vaccine is a top priority. In a pilot study, using a macaque model, we delivered HIV gp140 envelope glycoprotein and SIVmac239 P55 Gag and Nef antigens using heterologous prime/boost via the intranasal route with a soybean oil-based nanoemulsion (NE) adjuvant and through the intramuscular route with the AS01B adjuvant system to generate enhanced cell-mediated immunity. We used a NE adjuvant to promote gut-homing cell-mediated immunity and the AS01B system to enhance humoral immune responses. Following intrarectal challenge with SHIV 4MTF.tHy, vaccinated macaques acquired the virus but experienced lower viral loads in plasma (P=0.003) and CSF (P=0.001), and potent polyfunctional gag-specific (CD107a+, IFNγ, TNFα+) responses across diverse lymph nodes. Significant antibody-dependent complement deposition (ADCD) and antibody-dependent cellular phagocytosis (ADCP) responses were induced, and gut-microbiome crosstalk could be modulated and showing reduced SHIV-dysbiosis. Notably, vaccination preserved mucosal all-trans retinoic acid levels (atRA) (p<0.05). However, no significant differences were observed for antibody responses between vaccinated and unvaccinated macaques. In summary, the induced gut-homing properties by the NE adjuvant are effective at generating cell-mediated immunity and reducing viral set points and warrant further investigations as a mucosal adjuvant in HIV vaccine design.

**Importance:** Three major non-mucosal vaccine trials (RV144, HVTN702, and 706) failed to reduce HIV infection rates. Therefore, new approaches in developing a mucosal vaccine remain an effective strategy to attempt to control HIV infection. Coherent vaccine approaches against HIV were focused on immune correlates related to viral loads, persistent reservoirs, and antibody responses. As a proof-of-principle, we developed a vaccine regimen consisting of AS01B and an adjuvanted oil-in-water NE cleaved HIV clade C gp140 protein and non-cleaved Gag, and nef particles administered through intranasal, subcutaneous, and intramuscular routes, followed by intrarectal challenge with clade C SHIV. This vaccine elicited strong ADCD and ADCP responses, modulated immune-microbiome crosstalk, and reduced susceptibility to SHIV-infection-associated dysbiosis. Additionally, it preserved mucosal all trans retinoic acid (atRA) levels, suggesting a potential role for this approach in HIV vaccine development.

---

The Human Immunodeficiency Virus (HIV) epidemic remains a major public health concern and possesses a substantial economic burden. In 2023, the World Health Organization (WHO) reported approximately 39.9 million people living with HIV (PLWH), 630,000 deaths due to HIV-related causes, and 1.3 million new infections (1). Due to advanced antiretroviral therapy (ART), the life expectancies of PLWH have been extended to resemble those of uninfected individuals (2). However, despite prolonged longevity, the quality of life of persons exposed to ART is remarkably lowered due to drug toxicity, treatment fatigue, and increased occurrence of non-communicable diseases such as cardiometabolic abnormalities, neurological complications, and accelerated aging (3–5). ART has contributed to a reduced rate of new infections through pre-exposure prophylaxis and lowered transmission; however, an effective vaccine represents a more economical and efficient approach for reducing HIV cases and improving quality of life for PLWH.

HIV’s genetic diversity and ability to evade host immunologic clearance have impeded efforts to protect, control, and eradicate infection (6–9). Therapeutic vaccines are a promising HIV functional cure strategy developed to control viral replication, eliminate persistent reservoirs, and ultimately diminish the need for long-term ART (10). Currently, the RV144 trial is the only HIV vaccine clinical trial that has demonstrated limited protective efficacy (31.2%) associated with Env-specific antibodies and antibody-dependent cellular cytotoxicity (ADCC). For this trial, a canarypox vector ALVAC-HIV prime with a recombinant gp120 boost was utilized (11). Attempts to replicate and improve the effects seen in the RV144 trial have been unsuccessful. For instance, the HIV Vaccine Trials Network (HVTN) 702 trial, which used a bivalent gp120 MF59 adjuvant booster, was abruptly halted when HIV-1 infection was diagnosed in 5.1% of the vaccine recipients compared to 4.9% of placebo recipients (12). These HIV vaccine strategies focused primarily on establishing systemic immunity. However, since more than 90% of HIV infections are transmitted across mucosal tissues, primarily via sexual contact, efforts have been extended towards developing mucosal immunization strategies to enhance immunity at portals of entry (13).

Upon HIV encountering the mucosal barrier, a local infection is established, quickly spreading to lymphoid tissue and resulting in architectural destruction. During acute infection, there is an explosive replication of the virus, quantified by a high viral RNA level and typically a negative IgG/IgM-sensitive HIV-1 antibody test (14). However, the rapid virus replication, particularly in the gut-associated lymphoid tissue (GALT), leads to rapid depletion of GALT CD4+ T cells and later plateauing of viremia (14). Following peak viremia, the magnitude of viral load steadily drops towards a relatively stable viral set point. Individuals with a higher viral set point have been observed to have faster progression to acquire immunodeficiency syndrome (AIDS), thus making it a predictor of disease progression. As such, the magnitude of the viral set point has been delineated as a measure of vaccine efficacy (15–17). Several mucosal vaccine approaches have been explored to prevent HIV and simian immunodeficiency virus (SIV) transmission (18). These have included antigens ranging from recombinant bacterial and viral vectors, recombinant DNA, and purified proteins (19). Furthermore, use of envelope glycoprotein trimers has markedly improved in HIV-1 vaccine humoral responses. Recently, the HVTN 100 trial evaluated the immunogenicity profile of an HIV subtype C pox-protein vaccine regimen, which consisted of an ALVAC-HIV (vCP2438) prime and bivalent gp120/MF59 boost. Results from this study indicated that the protein boost improved the durability of the vaccine-induced immune responses (20). Additionally, a study conducted in rabbits and rhesus macaques reported the trimeric envelope cycP-gp120 immunogen to induce enhanced neutralizing activity and broad anti-V1V2 loop antibodies against HIV (21). As such, cleaved trimeric Env glycoproteins represent a superior option for inducing an immune response (22,23).

The primary barrier to mucosal immunization strategies is the lack of an effective and safe mucosal vaccine adjuvant (24). Mucosal vaccines, like their parenteral vaccine counterparts, must incorporate immune stimulatory motifs that induce pro-inflammatory cytokines and up-regulation of costimulatory molecules necessary for activating naïve lymphocytes. Currently, TLR agonists, cytokines including IL-12 and CCL28, attenuated bacterial enterotoxins like cholera toxin (CT) or dmLT, and bioadhesives such as chitosan have been explored as adjuvants for HIV vaccines (19). For example, a mucosal IL-4R antagonist HIV vaccination strategy implementing a SOSIP-gp140 booster was able to induce high-quality cytotoxic CD4+/CD8+ T cells and humoral responses in pigtail macaques (9). Another adjuvant that has demonstrated significant efficacy and minimal toxicity when employed intranasally is the oil-in-water NE adjuvant, which induced titers of antigen-specific serum IgG comparable to other standard vaccines, although this adjuvant has never been used in the context of HIV (25,26). To target humoral immune activation, the AS01B Adjuvant System was employed in a herpes zoster subunit vaccine (Shingrix) and proved to be highly efficacious (90.5%) in preventing herpes zoster during the ZOE-50 and ZOE-70 trials (27). This AS01B system in the Shingrix vaccine induced strong humoral immunity (28).

HIV infection is associated with dysregulation of the innate and adaptive immune responses at the gut-microbiota interface, including loss of CD4+ cells and impaired barrier function, each of which allows increased bacterial translocation and chronic immune activation (29,30). Due to this destruction or concurrently with it, the microbiome is significantly disrupted during HIV/SIV infection (31). The resultant dysbiosis is characterized by a depletion of key Firmicute taxa, including butyrate-producing bacteria in the family Lachnospiraceae (32,33). Even with effective viral suppression, the damage to the microbiome is never fully reconstituted, which has led to efforts to improve the gut microbiome composition (34,35). Therefore, in addition to reducing viral replication, an effective vaccine would ideally protect against dysbiosis, which further contributes to immune activation and associated tissue destruction.

Here, we present a proof-of-concept pilot study that sought to test whether the paired system of AS01B and NE adjuvanted with cleaved HIV clade C gp140 and non-cleaved Gag, and Nef antigens administered using IN primes and IM/subcutaneous boosts could produce robust protective humoral and cell-mediated immunity against simian-human immunodeficiency virus (SHIV).

## Results

### Study design for mucosal immune response and microbiome profiling

We included seven adult, female rhesus macaques (RM) in this pilot study. Four were immunized with non-cleaved HIV gp140 envelope glycoprotein and cleaved SIVmac239 P55 Gag and SIVmac239 Nef antigens delivered in mucosal adjuvants: prime/boost/boost NE Pure Soybean oil- and the AS01B system (Fig. 1). Sites of immunization doses and administration are outlined in Fig. 1B. Three unvaccinated RMs were considered a control group, and all monkeys were challenged intrarectally with low dose (10,000 TCID50) SHIV 4MTF.tHy. We collected blood, CSF, and rectal biopsies longitudinally for viral load quantification and immune response evaluation, feces for microbiome profiling, and isolated peripheral blood mononuclear cells (PBMCs) at necropsy for intracellular cytokine staining (ICS) analysis.

**Figure 1.**
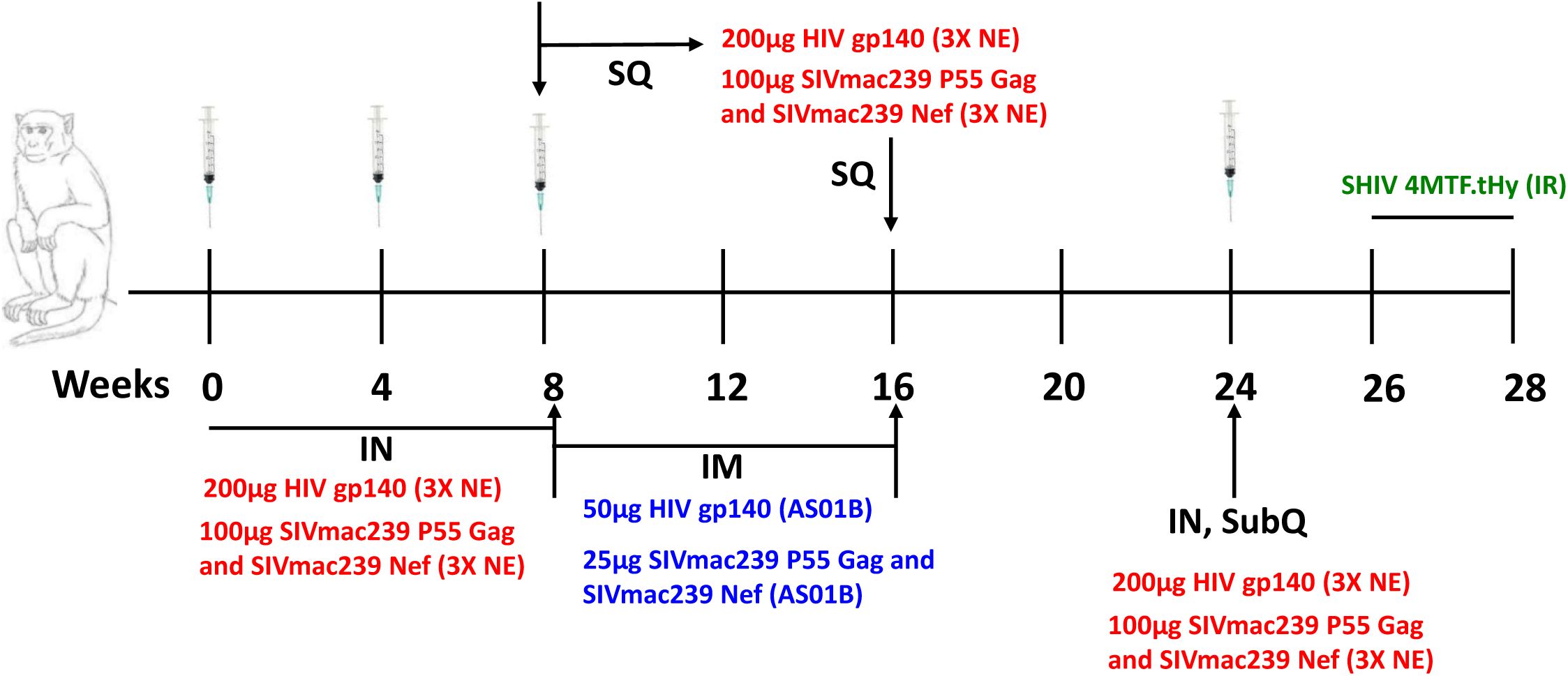
Immunization and challenge schema. (A) Study design. Four female rhesus macaques were immunized with HIV-1 Clade C env/gag-pol/nef antigens at weeks 0, 4, 8, 16, and 24. Two weeks after the last round of immunization, the four vaccinated monkeys along with three unvaccinated controls were challenged intrarectally with low-dose SHIV 4MTF.tHy. (B) Anatomical diagram illustrating the antigen dosages and sites of administration for intranasal (IN), intramuscular (IM), and subcutaneous (SubQ) routes.

### Vaccine-induced neutralization antibody responses

Neutralizing antibodies (nAbs) were measured using the TZM-bl assay against a tier 1A HIV-1 subtype C virus (MW965.26C) and SHIV virus (SHIV 4MTF.tHy) for plasma samples collected at weeks 0, 8, 12, and 18 during the vaccination period. Only week 18 plasma, at the peak immunity timepoint, showed weak neutralization against the tier 1A virus, although A13R040 also showed borderline levels at week 12 (Table 1). Only low titers were found against SHIV 4MTF.tHy were consistent across all time points, suggesting non-specificity. Since we did not see robust neutralization against either virus at any moment, we administered an additional IN/SubQ vaccine boost at week 24.

**Table 1.** Weak, nonspecific neutralizing antibodies generated by immunization in rhesus macaques. nAbs were measured by TZM-bl assay against a Tier 1A HIV and SHIV virus. Data are reported as ID_50_, i.e., serum dilution at which relative luminescence units (RLUs) were reduced by 50% compared to virus control wells, indicating the necessary value to inhibit 50% of viral infectivity. Values in black bold are considered positive for neutralizing antibody activity. The neutralization assays were performed until the 3rd immunization time point and currently planned to perform the 4th immunization time point.

| Animal ID | Bleed week | MW965.26 | SHIV<br>4MTF.tHY(P2)_A13T012 |
| --- | --- | --- | --- |
| A13R054 | 0 | <20 | 21 |
|  | 8 | <20 | <20 |
|  | 12 | <20 | 25 |
|  | 18 | 101 | 28 |
| A13R043 | 0 | <20 | 29 |
|  | 8 | <20 | 20 |
|  | 12 | <20 | 24 |
|  | 18 | 152 | 23 |
| A13R040 | 0 | <20 | 29 |
|  | 8 | <20 | 37 |
|  | 12 | 21 | 27 |
|  | 18 | 68 | 27 |
| A13R037 | 0 | <20 | 32 |
|  | 8 | <20 | 34 |
|  | 12 | <20 | 37 |
|  | 18 | 21 | 30 |
| CH01-31 | 0.5mg/mL positive<br>ctrl | 2.727 | nt |
| DH511.2_K3 | 0.5mg/mL positive<br>ctrl | <sup>1</sup>nt | 17.23 |
| VRC01 | 0.1mg/mL positive<br>ctrl | nt | 4.43 |

<sup>1</sup>nt: not tested

### Enhanced immune effector processes with vaccination

To assess the effect of our vaccine regimen on changes in antibody effector functions, we performed antibody-dependent complement deposition (ADCD) and antibody-dependent cellular phagocytosis (ADCP) analyses. In the vaccine group, a robust ADCD-inducing antibody response was observed at week 20, following the three IN primes and two subsequent IM/SubQ boosts (P < 0.05). This response was enhanced again at week 28, during peak viral load, coupled with a robust ADCP-inducing antibody response (P < 0.05), (Fig. 2A). At week 28, a borderline significant increment in ADCD was noted in vaccine recipients versus controls (P = 0.0571) while yielding a large effect size (d = 1.63), (Fig. 2B). Compared to baseline levels, ADCP-induced responses were elevated compared to baseline levels (P < 0.01) (Fig. 2C). Additionally, at week 28, comparing vaccinated and control rhesus macaques revealed a close to significant increase in ADCP levels (P = 0.0571) with a large effect size, (d = 1.74) (Fig. 2D).

**Figure 2.**
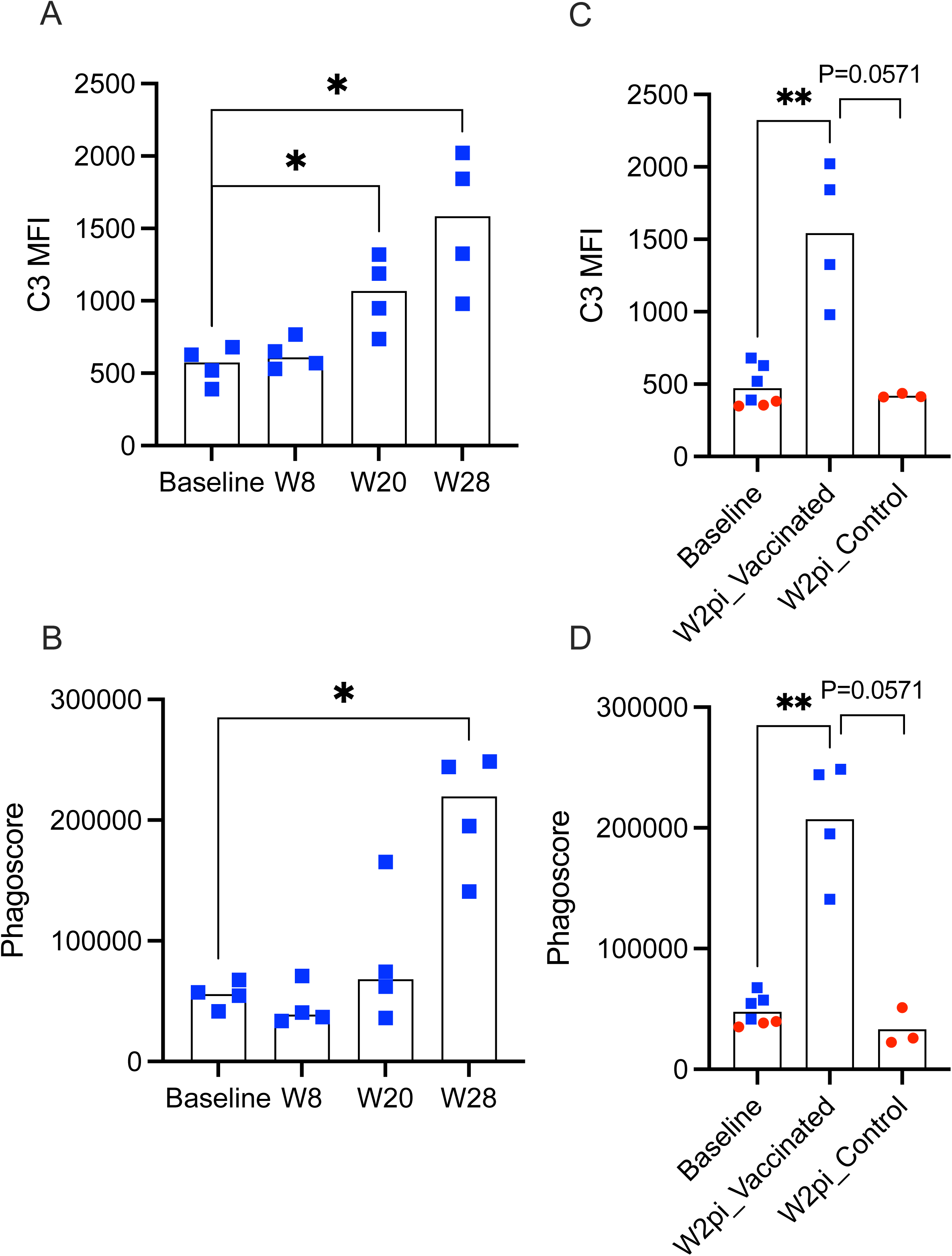
Vaccination enhances ADCD and ADCP induced immune responses. (A) ADCD responses quantified by C3 MFI scores and (B) ADCP responses quantified by phagoscore in vaccinated macaque plasma samples taken at baseline, and weeks 8, 20, and 28. Week 2pi corresponds to week 28. (C) Comparison of composite baseline ADCD or (D) baseline ADCP responses to week 2pi levels for vaccinated and control groups. Vaccinated macaques are denoted in blue, while control animals are in red.

**Vaccination lowers viral set point in plasma, CSF, and RB viral loads** as a readout for the efficacy of our vaccine regimen, we challenged the immunized RMs with the matched control unvaccinated animals as indicated above (Fig. 1A). All animals were followed longitudinally for 28 weeks post-infection to evaluate plasma, CSF, and cell-associated RB viral loads (VL) (Fig. 3A, C, E). Vaccination did not protect against the SHIV acquisition, given that all seven NHPs became infected after the first exposure. However, vaccinated monkeys had lower peak VLs than control monkeys during acute infection (week 2pi) (range: 7.1x10^6^ to 2.0x10^7^ vs 9.0x10^7^ to 3.7x10^8^ viral RNA copies/mL, respectively) and achieved lower viral set points such that one of the vaccinated animals had undetectable plasma VL at the steady state or viral set point; whereas two others had viral set point of <10^3^ SIV copies/mL (Fig. 3A). While not significant, likely due to our small sample size, the mean plasma viral set point at week 28pi (P=0.1) and mean CSF viral set point (P=0.1) for the immunized animals were lower compared to unvaccinated control set points (Fig. 3B, D). Considering CSF VLs, all the immunized animals except for the anomalous A13R037 had undetectable levels at the viral set point (week 28pi) (Fig. 3C). Additionally, tracking holistic groups mean VLs following infection, the immunized animals had significantly lower plasma (P=0.003**) and CSF (P=0.001**) VLs and a trend for lower RB VL compared to controls (Fig. 3B, D, F).

**Figure 3.**
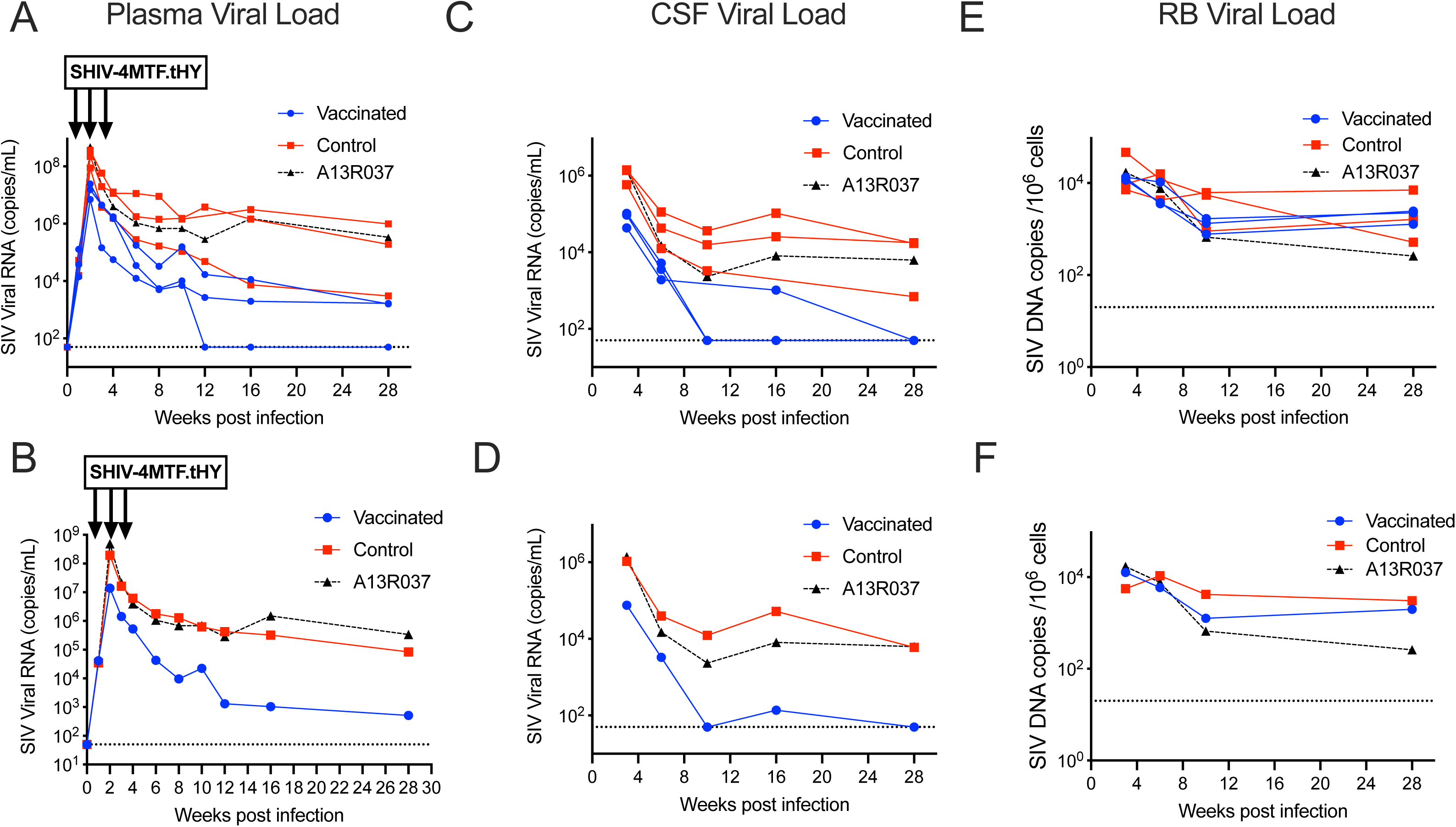
Vaccination reduced viral set point but did not protect NHPs from SHIV challenge. Rhesus macaques (n=4 in vaccine group; n=3 in control group) were challenged intrarectally weekly using a low dose (10,000 TCID_50_) of SHIV 4MTF.tHy. All animals received two additional challenges after the infective exposure. (A) Longitudinal plasma viral loads of individual animals after infection as quantified by RT-qPCR. (B) Geometric means of plasma viral loads (P=0.003*) (C) Longitudinal CSF viral loads of individual animals after infection as quantified by RT-qPCR. (D) Geometric means of CSF viral loads (P=0.001*) (E) Longitudinal rectal biopsy viral loads after infection as quantified by ddPCR. (F) Geometric means of RB viral loads. The dashed lines represent the limit of detection for the assay (50 copies/mL for qRT-PCR and 20 copies/mL for ddPCR). Vaccinated monkeys are denoted in blue, control monkeys in red, and the anomalous vaccinated monkey (A13R037) is in black.

### Cell-associated SIV DNA in the reproductive, gut, and lymphatic tissues

At necropsy, viral DNA copies were evaluated in reproductive, lymphatic, and gut tissues to understand how our mucosal vaccine regimen may have influenced the dissemination of the virus in the different compartments. We measured cell-associated viral DNA copies in the vagina, uterus, cervix, ovaries, and fallopian tubes within the reproductive tract. While the vagina and uterus failed to show the presence of any virus, we detected cell-associated viral DNA in the tissues of the cervix, ovaries, and fallopian tubes. Comparing the vaccinated and control monkeys, there are minimal differences within the cervix and fallopian tubes; however, the ovaries showed lower levels, though nonsignificant, of viral DNA for the vaccinated animals (P=0.4) (Fig. S1A). Further, within the lymphatic tissues, we measured viral DNA in the spleen, mesenteric, and inguinal LN tissues. There was a nonsignificant trend for lower viral levels in the mesenteric LN (P=0.7), inguinal LN (P=0.1), and spleen (P=0.1), suggesting that our vaccine generated effective anti-viral systemic immunity within the lymphatic system (Fig. S1B). To dissect a potential mechanism of action and understand mucosal immunity, we also quantified cell-associated viral DNA levels in specific layers of different regions of intestinal tissues. We considered the epithelium and lamina propria (Epi LP) as well as muscularis and serosa (Musc Sero) from the duodenum (Duo), ileum, and ascending colon (AC). Across these regions, levels were comparable between the vaccinated and control animals; the Musc Sero of the ileum (P=0.4) and Epi LP of the AC (P=0.1) showed lower cell-associated viral DNA levels for immunized RMs (Fig. S1C).

### Vaccine-mediated polyfunctional CD4+ and CD8+ T cell responses

The effect of exposure to the selected prime-boost vaccine combination on the induction of potent gag-specific responses was evaluated using peripheral blood mononuclear cells (PBMCs) and diverse lymph nodes (axillary, cervical, colonic, inguinal, and mesenteric) as denoted in Fig. S2 and 3. Borderline increments in gag-specific CD107a+ responses in both CD4+ and CD8+ T cells were noted across all the studied compartments (Fig. 4A and 4B, respectively). On the other hand, vaccinated rhesus macaques had borderline increments in CD4+ gag specific interferon-gamma (IFNγ) responses in only the axillary lymph nodes (Fig. 4C). Interestingly, significantly robust CD8+ gag IFNγ responses, notably within the axillary and inguinal lymph nodes, were observed in vaccinated rhesus macaques (all P < 0.05), (Fig. 4D). No gag specific CD4+ Tumor Necrosis Factor-alpha (TNFα) + responses were noted across all studied tissue compartments, (Fig. 4E). Alternatively, significant CD8+ TNF α+ responses were noted within the inguinal lymph nodes of vaccine exposed rhesus macaques. In addition, borderline increments of CD8+ TNF α+ responses were noted in axillary and cervical lymph nodes.

**Figure 4.**
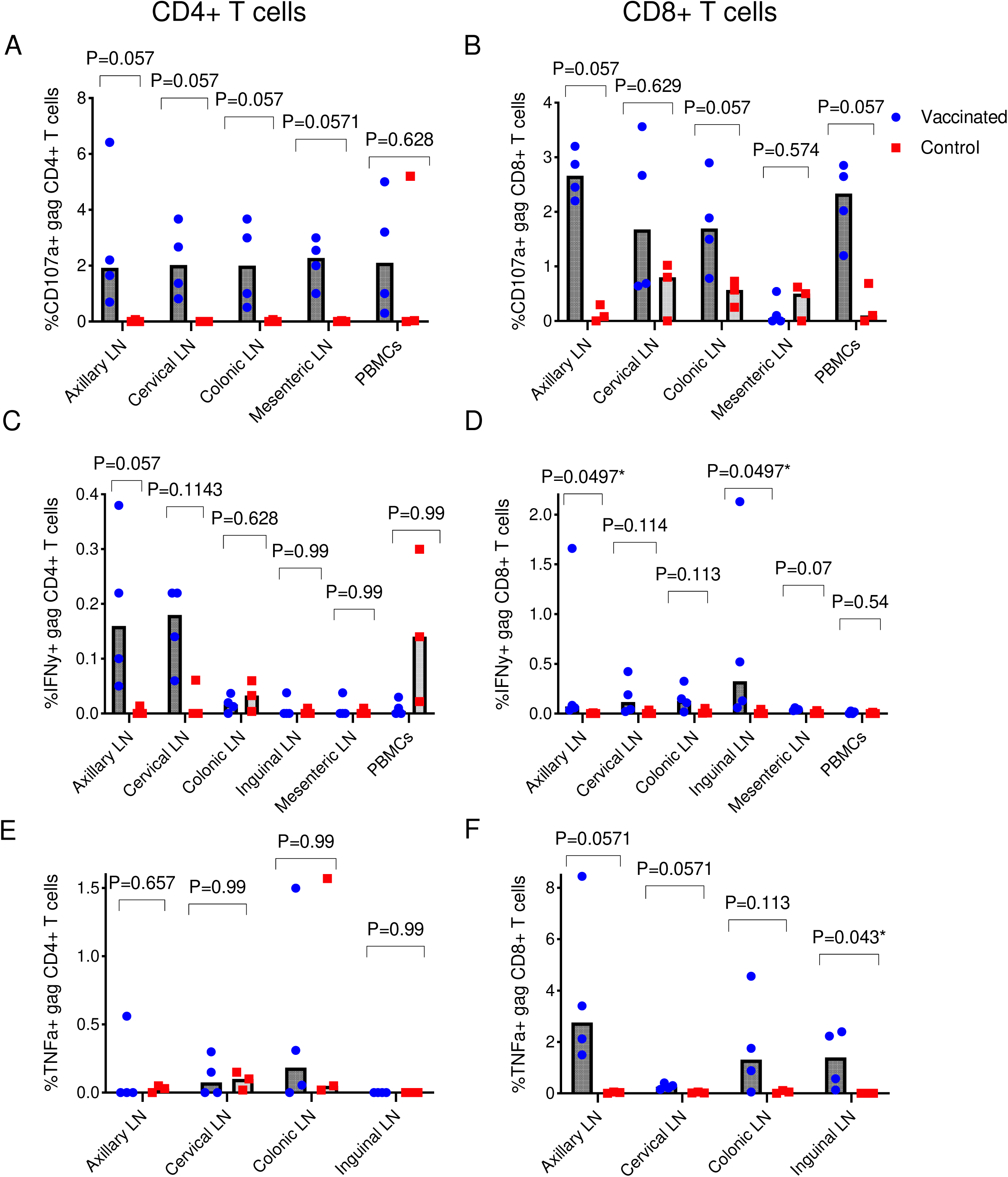
Vaccination induces gag specific T cell responses in diverse tissue niches. (PBMCs, and mesenteric, colonic, inguinal, cervical, and axillary lymph nodes). Necropsy timepoint comparisons of either (A) CD4+ and (B) CD8+ gag specific CD107a+ responses in vaccinated versus non-vaccinated rhesus macaques. Changes in (C) CD4+ and (D) CD8+ gag specific IFN γ levels in diverse lymph nodes and PBMCs of vaccine versus control rhesus macaques. Alteration in (E) CD4+ and (F) CD8+ gag specific TNFα levels of diverse lymph nodes and PBMCs of vaccinated versus control rhesus macaques.

### Mucosal vaccination facilitates the preservation of intestinal retinoid metabolism

Mucosal damage leads to significant disruption to all-trans retinoic acid (atRA) synthesis in the gut (36). This pattern is also seen during SIV infection of Asian macaques, a finding consistent with current models suggesting that gut damage is a primary driver of HIV/SIV pathogenesis (37,38). Previous research has determined that NE formulations facilitate increased atRA synthesis in the gut by activating RALDH activity in dendritic cells, thereby modulating lymphocyte differentiation, activation, and homing capabilities (38). To determine whether retinoids may be contributing to mucosal vaccination efficacy in our study, the longitudinal levels of atRA and its precursors retinol (ROL) and retinyl esters (RE) were analyzed in the plasma by mass spectrometry. No significant differences were found in plasma atRA, although there was a trend in higher atRA by the time of necropsy (Fig. 5A). Similarly, no differences were found in atRA precursors ROL and RE at any time point (Fig. S4A, C).

**Figure 5.**
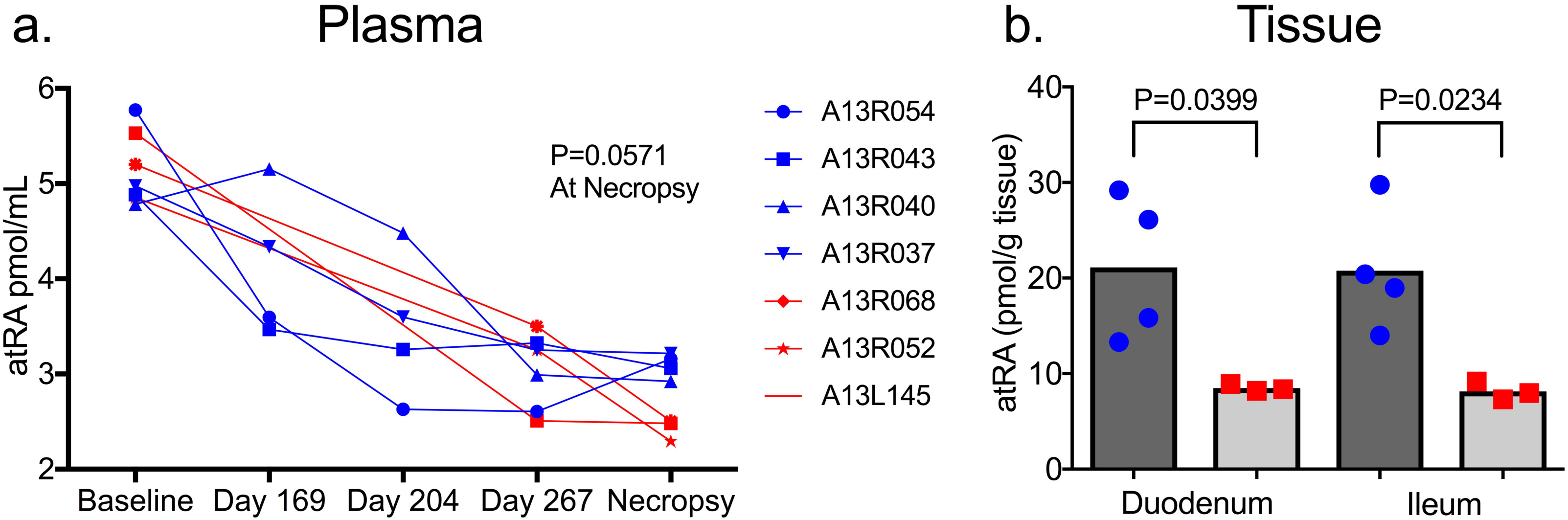
Vaccination facilitates elevated retinoic acid synthesis in mucosal tissue. (A) Comparisons of plasma all-trans retinoic acid following SHIV challenge. Tissue all-trans retinoic acid for both the (B) duodenum and ileum of the small intestines.

Although minimal changes were found in the plasma, we presumed atRA would be elevated in the mucosal tissue following our vaccination. Gut tissue from the duodenum and ileum was collected at necropsy, and retinoid concentrations were similarly determined. Vaccination was associated with significantly higher atRA in both the duodenum (P=0.04) and the ileum (P=0.02) (Fig. 5b). Additionally, there was a lower ROL and RE trend in the duodenum of the vaccinated group at necropsy (Fig. S4B, D). This finding suggests that more atRA precursors are utilized as substrate for atRA synthesis following vaccination, consistent with previous data for similar NE formulations, indicating that it increases mucosal RALDH activity (39). Additionally, ROL and RE trend higher in the duodenum than in the ileum. To the best of our knowledge, this data is the first to validate a previous finding in a large animal model that mouse retinoid concentrations trend lower from proximal to distal tissue in the gastrointestinal tract (40).

### Vaccination Reduces Acute SHIV Infection-Associated Dysbiosis

During acute and chronic HIV infections, distinct dysbiosis patterns are characterized by the expansion of Bacteroidetes and loss of Firmicutes, particularly butyrate-producing members of Lachnospiraceae (32,41). In addition to protection from viremia and lymphoid cell destruction, an ideal mucosal vaccine strategy would assist in maintaining microbial eubiosis. To monitor gut microbiome dynamics, we performed 16S rRNA sequencing and found dysbiosis markers in the control group but not in the vaccinated group during acute infection. This dysbiosis was characterized by the expansion of bacterial families Muribaculaceae and Fibrobacteraceae (Fig. 6). Other less abundant families were also expanded at Day 21pi/week 3pi, including Oligosphaeraceae, Erysipelothrichaceae, WCHV1_41_unclassified, and Bradymonadales_unclassified (Fig. 6A). Additionally, the family Lachnospiraceae was depleted during acute infection. Of note, the major butyrate-producing member of Lachnospiraceae *Roseburia* showed a particularly large decrease during acute infection (Fig. 6B).

**Figure 6.**
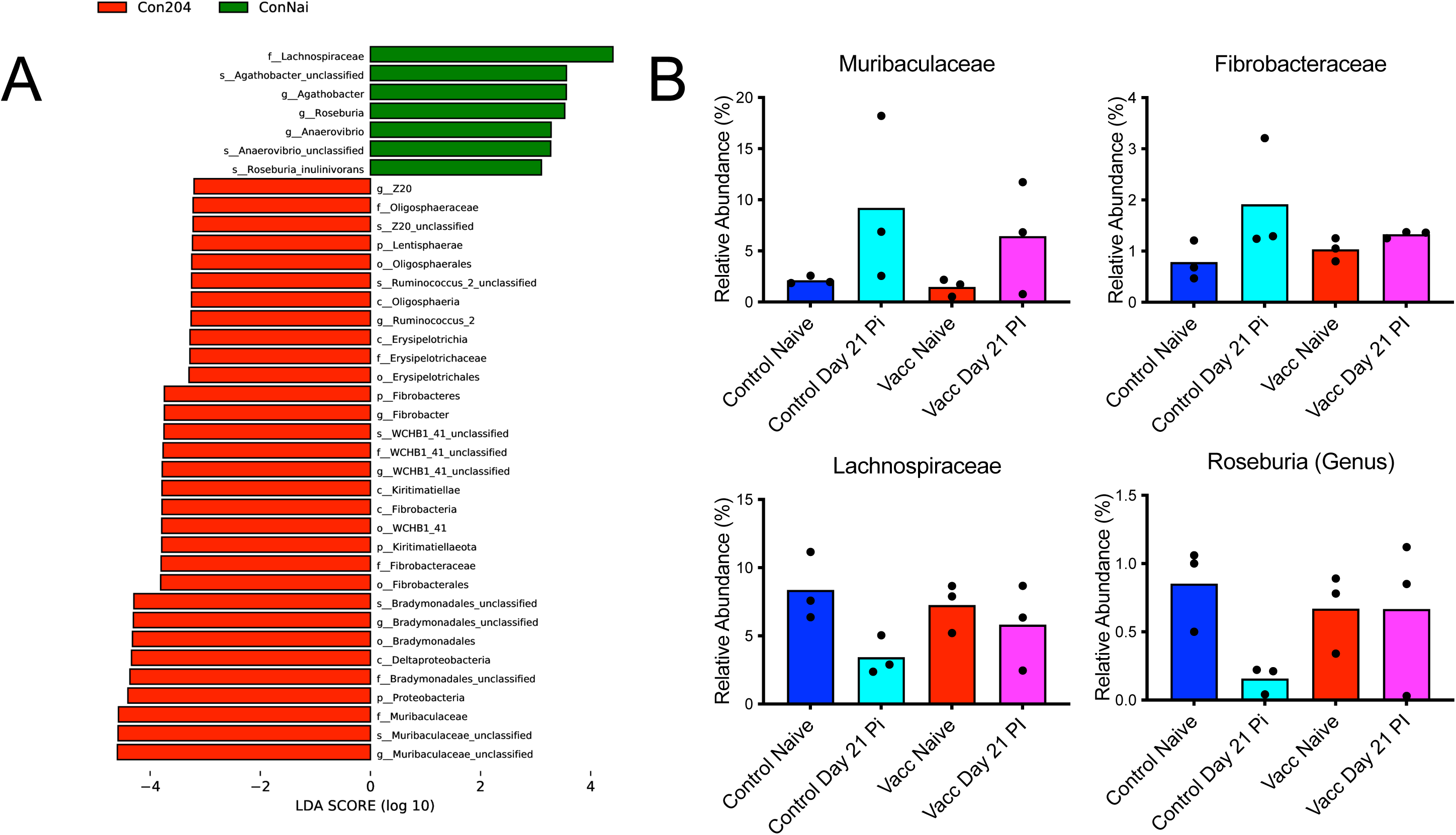
Vaccination protects against acute infection-associated dysbiosis. (A) LEfSE analysis was performed for all timepoints for each group. Only control Day 204 (Con204) (Day 21pi/week 3pi) resulted in significant changes compared with baseline (ConNai). This dysbiosis was characterized by (B) expansion of Muribaculaceae and Fibrobacteraceae families and loss of Lachnospiraceae, particularly the butyrate-producing genus *Roseburia*.

Certain bacteria are adept at modulating host mucosal immunity, and *Helicobacter* specifically has been tied to vaccine efficacy (42,43). Following vaccination, but before infection, the anomalous vaccinated monkey (A13R037) had a significant expansion of *Helicobacter* (Fig. S5A). In the vaccinated monkeys, the relative abundance of *Helicobacter* at this timepoint was positively associated with acute infection viral loads at Day 21pi/week 3pi (r=0.9921, p=0.0079), suggesting that *Helicobacter* specifically may be a factor in A13R037’s breakthrough infection (Fig. S5B).

### Antibody responses in CVL fluid and plasma

We evaluated HIV clade C envelope gp140-specific IgG (plasma) and IgA (CVL and plasma), as well as total IgG and IgA (CVL) antibody responses. Our results showed that only at week 28 (D28pi/D211), the vaccinated group presented a higher gp140 HIV-specific IgA titer compared to the control RM’s. Likewise, the gp140 HIV specific IgG titer was lower at week 32 (D56pi/D239) and week 40 (D112pi/D295) in the vaccinated group compared to the control RM’s. We did not observe any significant differences in the CVL antibody responses between the vaccinated and control RMs at different time points (Fig. S6).

## Discussion

HIV pathogenesis has been linked to the loss of peripheral CD4+ T cells during acute infection, chronic immune activation, and compromised integrity of the intestinal barrier, allowing microbial translocation, thus exacerbating the dysfunction. In the present study, we show a vaccination strategy that targets retinoic acid signaling in the gut, typically lost during HIV/SIV pathogenesis, and may play a substantial role in facilitating improved viral control. This strategy improved both the polyfunctionality of CD8+ T cells at inguinal lymph nodes and the phagoscore in ADCD and ADCP function. Additionally, this mucosal vaccine protected butyrate-producing bacteria typically depleted during acute infection (33). Butyrate has a pleiotropic role in the gut, acting as an energy source for colonocytes, a histone deacetylase inhibitor, and a GPR109A agonist regulating macrophages in the mucosa (44,45). Previously, we have also found that fecal butyrate levels were associated with expression of colonic ALDH1A2 and ALDH1A3, two genes involved in atRA synthesis, during SIV infection in macaques (38). In the present study, mucosal atRA was preserved at the mucosal sites, but not systemically, suggesting that the aim of improving mucosal immune response was effective and highly localized. It may be dependent on improved retinoid signaling in the gut that is lost during HIV/SIV infection and never recovered, even during cART.

The current study proves that the paired adjuvants AS01B and NE with HIV-1 env-gag-nef antigens are immunogenic and safe. Our vaccination strategy generated minimal CD4+ and CD8+ T cell virus-specific responses within the periphery and lymphoid tissues, provided robust ADCD and ADCP responses, activated the retinaldehyde dehydrogenase (RALDH) pathway, and modulated immune-microbiome crosstalk. Pure soybean oil NE adjuvant was chosen to achieve gut homing through its effect on RALDH (39). However, the ASO1B adjuvant system was added to stimulate broadly neutralizing antibodies and cell-mediated immunity in the Shingrix vaccine with 95% effectiveness (27). However, the proof of the concept of “gut homing” was achieved, as evidenced by the preservation of the atRA and gut microbiome in the vaccinated animals, and may have contributed to lowering viral set points in the vaccinated animals. Even though the exact mechanism by which elite controllers and long-term non-progressors maintain low viral load is not clear (46,47), the viral set point control achieved in our vaccinated macaques may provide some leads to investigate in these cohorts. What role the ASO1 B system played in the low viral set point is not clear. This would require a study using both adjuvants separately. In this study, we did not look at the CD4+ and Th17 T cell levels in the GALT and the intestinal mucosa; this would be a valuable marker and the effectiveness of the vaccine in future studies.

Most notably, following intranasal primes and subsequent intramuscular/subcutaneous boosts of our HIV vaccine and then challenge with SHIV, rhesus macaques exhibited lower viral set points. Such that the viral burden in both plasma and CSF was significantly lower than their control counterparts. While all animals within the vaccinated group became infected after a single challenge with the virus, they did exhibit better virus control. Recently, a lower viral set point has been treated as a readout for vaccine efficacy (48). Further, an antiretroviral therapy interruption (ATI) study that involved the administration of the HIV vaccine candidate, Vacc-4x, demonstrated a significant reduction in viral load set point upon ATI. A follow-up to this investigated the maintenance of the Vacc-4x effect by re-boosting eligible participants and observed promising results for Vacc-4x as a therapeutic vaccine in comparably lowered viral set point compared to pre-ART levels (49). As expected, the vaccine-related reduction of viral loads was accompanied by improved virus-specific cellular immune responses. Like others, we noted that vaccine administration induced gag-specific CD4+ and CD8+ T cell responses (CD107a+, IFNγ+, and TNFα+), which were associated with vaccine-mediated control of viremia (50–52).

Beyond induction of cellular immunity, we also noted that our vaccine regimen did not induce the same along with neutralizing antibody responses against Tier 1/2 viruses, or the SHIV used for the challenge. This data suggests an alternative mechanism for improved virologic control as previously reported (53,54) and might destroy virus-infected cells through Fc-mediated functions such as ADCD or ADCP (55–57). Our study showed strong ADCD or ADCP responses at week 20 and during acute infection at peak viral load (week 28), coupled with augmented phagocytosis, which provides further evidence that our vaccine regimen directly targets infected cells through diverse mechanisms. (58,59). Like the RV144 vaccination regimen, enhanced Fc-mediated effector functions could highlight improved vaccine-induced B cell priming, resulting in augmented downstream antibody efficacy and function (60).

Additionally, the adjuvant system played a significant role in the increased efficacy of our vaccine. The NE adjuvant has been implicated in influenza vaccination and allergy treatments, but to date, it has not been exploited in an HIV vaccine (61, 62). A recent study demonstrated that IN immunization with NE indirectly activated retinaldehyde dehydrogenase activity in dendritic cells and led to an expression of alpha-4-beta-7 (α4β7) integrin and CCR9 gut-homing receptors on T cells, culminating in robust mucosal cell-mediated immunity (46). This is supported by enhanced levels of atRA in the gut tissues of vaccine recipients as compared to controls. For the first time, our study demonstrated that the NE adjuvant could be employed to enhance mucosal immune responses in the context of HIV because our study also demonstrated an increase in gut-specific atRA levels in vaccinated animals. In contrast, atRA levels in the periphery remained unchanged compared to controls.

In addition to lifelong infection, HIV alters the gut microbial composition, even during ART suppression, by a poorly defined mechanism (31). Because of its role in immune modulation and use as a biomarker, 16S rRNA sequencing was performed to understand the microbiome composition. Control animals experienced acute dysbiosis characterized by increases in Muribaculaceae and Fibrobacteraceae, and decreases in Lachnospiraceae, most notably *Roseburia*, a butyrate-producing genus associated with lower levels of immune activation and microbial translocation markers during HIV infection, each of which contributes significantly to accelerated biological aging, even during ART (41). We have also reported a role for *Roseburia* mucosal immunotherapies during SIV infection, including its abundance being associated with lamina propria macrophage maturation and PD-1 expression on CD4+ central memory T cells, which contributes substantially to the viral reservoir (63, 33). The 16S rRNA sequencing in the present study also demonstrated an association between acute viral load and the relative abundance of *Helicobacter* following vaccination. We hypothesize that the expansion of *Helicobacter* seen in A13R037 may partially account for its breakthrough infection. Sequence alignment was unable to determine the predominant *Helicobacter* species in our study. However, previous work has suggested that the relative dominance of *Helicobacter* in rhesus macaque microbiomes compared with humans may impact translational studies (64). *Helicobacter* spp. is a potent immune modulator able to block activation, inhibit phagocytosis, and either increase or decrease gut trafficking in their respective hosts (65). Additionally, *Helicobacter pylori* is associated with the suppression of atRA synthesis in the gastric mucosa (66). Rhesus macaque-specific *Helicobacter* spp. have been implicated in colitis, but how they impact gut immune trafficking and function in the context of vaccination remains poorly understood (67).

The study presented here provides evidence that pairing AS01B with an NE adjuvant generates a cellular immune response, reduces peripheral and CNS viral loads, and protects the microbiome from acute dysbiosis and significant depletion of retinoic acid signaling from the small intestines. Although sterile immunization was not achieved, this strategy offers proof of principle that such mucosal vaccination strategies may play a key role in efforts to reduce HIV transmission and the development of future therapeutic vaccination strategies for people already living with HIV. For the first time, we demonstrated that the gut-homing properties of the NE adjuvant are effective at generating cell-mediated immunity in the context of HIV and may be pursued as a mucosal adjuvant in the future of HIV vaccine designs.

## Materials and Methods

### Animals and Ethics Statement

Seven outbred, pathogen-free, adult female rhesus macaques (*Macaca mulatta*) of Indian origin with a mean age of 7 years were used in this study (**Table 2**). Macaques were socially housed in pairs in a temperature-controlled (72℉) indoor climate with a 12-hour light/dark cycle. Housing followed regulations set by the Animal Welfare Act and Guide for the Care and Use of Laboratory Animals in nonhuman primate facilities at the Department of Comparative Medicine, University of Nebraska Medical Center (UNMC), Omaha, NE, USA. Animals were fed commercial primate chow (Teklad; 2055) supplemented with fruits and vegetables daily and provided R/O filtered municipal water ad libitum. Environmental enrichment was provided daily through toys, videos, and treats. Additionally, RMs were observed daily for signs of stress or disease by veterinary personnel and animal care staff. After the study, RMs were humanely euthanized, according to the American Veterinary Medical Association, by using a high dose of ketamine-xylazine followed by the opening of the thoracic cavity and perfusion/exsanguination. This study, under protocol number 18-108-08-FC was reviewed and approved by the UNMC Institutional Animal Care and Use Committee (IACUC) and the Institutional Biosafety Committee (IBC). UNMC has been accredited by the Association for Assessment and Accreditation of Laboratory Animal Care International.

**Table 2.** Identifiers and information about rhesus macaques used in this study.

| Serial no. | Animal ID | Age (yrs) | Treatment group | Plasma Viral load (copies/mL) |  | CSF Viral Load (copies/mL) |  |
| --- | --- | --- | --- | --- | --- | --- | --- |
|  |  |  |  | Peak | At necropsy | Peak | At necropsy |
| 1 | A13R054 | 7 | Vaccinated | 7.1E+06 | 1.6E+03 | 4.3E+04 | <LOD |
| 2 | A13R043 | 7 | Vaccinated | 2.5E+07 | <LOD<sup>b</sup> | 1.0E+05 | <LOD |
| 3 | A13R040 | 7 | Vaccinated | 1.5E+07 | 1.6E+03 | 9.4E+04 | <LOD |
| 4 | A13R037 | 7 | Vaccinated | 4.9E+08 | 3.4E+05 | 1.4E+06 | 6.3E+03 |
| 5 | A13R068 | 7 | Control | 9.0E+07 | 3.0E+03 | 5.8E+05 | 7.0E+02 |
| 6 | A13R052 | 7 | Control | 3.7E+08 | 9.9E+05 | 1.4E+06 | 1.7E+04 |
| 7 | A13L145 | 7 | Control | 2.3E+08 | 1.9E+05 | 1.4E+06 | 1.8E+04 |

<sup>a</sup>ID, identifier. All animals were female
<sup>b</sup>LOD, limit of detection

### Study design

The overall design of the study is outlined in schematic form in Fig. 1. The experiments conducted here were designed to evaluate whether an HIV vaccine combining mucosal adjuvants with immunogenic envelope glycoprotein of HIV clade C can elicit mucosal immunity in SHIV-infected RMs. We included seven RMs in the study that were randomly divided into two experimental groups. The first group (n=4) was vaccinated and the second (n=3) was the control group. We longitudinally collected blood, CSF, feces, and rectal biopsies (RB) before and during the vaccination period, as well as throughout the infection (Fig. 1A).

### Immunization and challenge

The immunization and challenge schedules are outlined in Fig. 1. On weeks 0, 4, and 8, the RMs in the vaccinated group (n=4) were primed with two intranasal (IN) injections. IN injections were made using an intranasal mucosal atomization device (Teleflex Medical, #MAD100); the first was administered into one nostril consisting of 200 μg HIV gp140 envelope glycoprotein (CH505TFchim.6R.SOSIP) (Duke University) adjuvanted with 3xNE Pure Soybean oil-nano Emulsion (Dr. Baker, University of Michigan) (1:2), and the second was administered into the contralateral nostril consisting of 100 μg SIVmac239 P55 Gag and 100 μg SIVmac239 Nef antigens (NHP Resources, University of Lafayette (Dr. Villinger)) in 3xNE adjuvant (1:2). On weeks 8 and 16, the RMs in the vaccinated group received two intramuscular (IM) injections into opposite quadriceps muscles. The first was comprised of 50 μg HIV gp140 envelope glycoprotein in AS01B Shingrix adjuvant, and the second consisted of 25 μg SIVmac239 P55 Gag and 25 μg SIVmac239 Nef antigens adjuvanted with AS01B Shingrix. In weeks 8 and 16, RMs also received two subcutaneous (SubQ) injections into the contralateral skin layers of the abdominal wall quadrant proximal to the mid inguinal ligament area. SubQ mixtures were identical to IN. Additionally, on week 24, RMs received a boost administered intranasally and subcutaneously as performed before. At week 26, all seven RMs were challenged weekly for three total visits by the intrarectal (IR) route with 10,000 TCID_50_ of our SHIV 4MTF.tHy viral stock (68). This virus stock was generated by transfecting the SHIV plasmid into 293T cells, and the resulting virus supernatant was used to infect macaque PBMCs. A large prep of virus stock was generated and titered in TZM-bl cells.

### Neutralization assay

Plasma samples for all animals in the vaccinated group (n=4) were collected at baseline (naïve), and weeks 8, 12, and 18 during the immunization period for measuring neutralization. The TZM-bl neutralization assay was carried out as described previously (69). The IgG concentration required to achieve 50% neutralization was reported as the 50% inhibitory concentration (ID50) and was calculated to concerning virus-only wells. Values were calculated using the following formula: [(value for virus only minus value for cells) minus (value for serum minus value for cells only)] divided by (value for virus minus value for cells only).

### SHIV Plasma and CSF viral load quantification

*Gag*-targeted quantitative reverse transcription-PCR (qRT-PCR) was employed to measure SHIV RNA concentration in plasma and CSF samples. Blood was collected from the femoral vein in K2-EDTA vacutainer tubes (Becton), and within 2 hours of collection, the plasma was separated by centrifugation at 1200 rpm for 20 minutes. CSF was collected by lumbar puncture, centrifuged at 2000 rpm for 10 minutes, and then the supernatant was separated for RNA extraction. Following the manufacturer’s instructions, the QIAamp viral RNA Minikit (Qiagen, 52906) was used to isolate RNA from 140 μL of plasma and CSF samples. The TaqMan RNA-to-Ct 1-Step kit (Thermo Fisher Scientific, 4392938) and Applied Biosystems QuantStudio 3 real-time PCR system (Applied Biosystems) were used to quantify SIV Gag RNA by qRT-PCR. The following primers and probes were used: SIVGAGF, 5′-GTCTGCGTCATCTGGTGCATTC-3′; SIVGAGR, 5′-CACTAGGTGTCTCTGCACTATCTGTTTTG-3′; and SIVP, 5′-/6-carboxyfluorescein (FAM)/CTTCCTCAG/ZEN/TGTGTTTCACTTTCTCTTCTGCG/3IABkFQ-3′.

### Isolation of DNA from RB frozen tissues

For DNA isolation from frozen rectal biopsy tissues, the DNEasy Blood and Tissue kit (QIAGEN, #69506) was used. In brief, a single pinch biopsy was cut into pieces (<10 mg) and placed into a 1.5-mL microcentrifuge tube with 180 μL Buffer ATL and 20 μL Proteinase K. Samples were incubated at 56°C for 3 five-minute periods with thorough vortexing in between. Following homogenization, 200 μL of Buffer AL was added, and samples were incubated at 56°C for 10 minutes. DNA was then isolated by the manufacturer’s protocol and eluted in 50 μL of Buffer AE. DNA concentrations were measured using a SimpliNano spectrophotometer (GE Healthcare Bio-Sciences Corp).

### Isolation of DNA/RNA from frozen tissues

DNA/RNA was isolated from frozen tissues using the AllPrep DNA/RNA minikit (Qiagen, 80204). Approximately 30mg of frozen tissue and a 5-mm stainless steel bead (Qiagen, 69989) were added to a 2-ml flat-bottom tube on dry ice and equilibrated for 20 minutes. Next, 600 μL of buffer RLT plus supplemented with 2-mercaptoethanol (ßME) (10 μL/mL of RLT plus) was added, followed by homogenization using a TissueLyser LT (Qiagen, #69980) set at 50 oscillations/sec for 8 minutes. After checking for homogeneity, the lysate was transferred to a QIAshredder (Qiagen, 79656) and then DNA/RNA was isolated per the manufacturer’s instructions. DNA was eluted in 50 μL of EB buffer, while RNA was eluted in 50 μL of RNase-free water. A SimpliNano spectrophotometer was used to measure DNA/RNA concentrations.

### SHIV cell-associated DNA quantification

Total cell-associated SHIV DNA was measured by ddPCR using 1 μg DNA, 2X ddPCR Supermix for probes (no dUTP) (Bio-Rad, 1863024), and the same primer-probe set used to quantify SIV plasma viral load. The Bio-Rad QX200 AutoDG digital droplet PCR system was used for the procedure. To do so, 22 μL of reaction mix was used for droplet generation, then the ddPCR plate containing the emulsified samples was heat sealed with foil (Bio-Rad, 181-4040) and transferred to a C1000 Touch thermal cycler (Bio-Rad). Following thermal cycling, the ddPCR plate was moved to a QX200 droplet reader (Bio-Rad) for fluorescence measurement and droplet count. QuantaSoft software was used to separate positive droplets with amplified products from negative droplets by applying a fluorescence amplitude threshold and to measure the absolute quantity of DNA per sample (copies/mL). To quantify the input cell numbers, a parallel ddPCR reaction for the rhesus macaque host gene, RPP30, was carried out.

### Intracellular Cytokine Staining (ICS) of gag-specific T cells across diverse tissue compartments (peripheral blood and various lymph nodes)

Fresh PBMCs were isolated from rhesus macaque peripheral blood using discontinuous density gradient centrifugation on Percoll. Next, a brief 6h rest was followed by stimulation with SIV mac239 gag peptide mixes spanning 15-mers with 11-aa overlap (ARP-6204) supplied by the NIH HIV Reagent Program, Division of AIDS, NIAID, NIH: Peptide Array, Simian Immunodeficiency Virus (SIV)mac239 Gag Protein, ARP-6204, contributed by DAIDS/NIAID) and PMA (Phorbol myristate acetate) /ionomycin (20 ng/ml and 0.5 μg/ml). Monensin was added as an intracellular transport inhibitor to stop the release of proteins into the supernatant. After 6 hours of stimulation, a 20mM EDTA solution was added to stop the reaction. Subsequent washes were then performed, and live/ dead exclusion was performed using Zombie Aqua dye. Further washes and later fixation with 2% paraformaldehyde (PFA) were then done. Next, BD perm wash buffer was used for permeabilization, followed by the addition of a freshly prepared cocktail of all antibodies. Following a 30-minute incubation and later washes, the acquisition was then done using a Becton Dickson Fortessa X450 flow cytometer. Data analysis was then carried out using Flowjo version 10.6 (Trees Star Inc., Ashland, Oregon, USA).

### Antibody-Dependent Complement Deposition (ADCD) Assay

ADCD was assessed using a flow cytometry-based complement-fixing assay (70). Briefly, 10 x 106 HUT78 cells were pulsed with 20µg recombinant HIV-1 gp120 from strain CN54 (Acro Biosystems) for 20 min at 37°C. Excess, unbound gp120 was removed by washing cells twice with 1% PBS-BSA buffer. Heat-inactivated monkey plasma samples diluted at 1:20 were added to the antigen-pulsed cells and incubated for another 30 min at 37°C. Freshly resuspended lyophilized guinea pig complement (Cedarlane) diluted 1:20 with veronal buffer, 0.1% gelatin with calcium and magnesium (Boston BioProducts) were added to the cells for 2 h at 37°C. Following wash with 1X PBS, cells were assessed for complement deposition by staining with goat anti-guinea pig C3-FITC (MP biomedicals). After fixing, cells were analyzed by flow cytometry, and ADCD is reported as the MFI of FITC+ cells

### Antibody-Dependent Cellular Phagocytosis Assay (ADCP)

ADCP was measured using a flow cytometry-based phagocytosis assay (71). Biotinylated recombinant HIV-1 gp120 from strain CN54 (Acro Biosystems) was combined with fluorescent NeutrAvidin beads (Life Technologies) at 1µg protein per µl bead and incubated overnight at 4°C. Beads were washed twice with 0.1% PBS-BSA to remove excess, unconjugated antigens. gp120-coated beads were resuspended in a final volume of 1 ml in 0.1% PBS-BSA. 10 μl bead suspension was added into each well of a round-bottom 96-well culture plate, after which heat-inactivated plasma samples were added (1:500) and incubated for 2 h at 37°C. 5 x 1p0^4^ THP-1 cells in 200 µl growth medium were added to each well and incubated overnight at 37°C. The following day, 100 μl of supernatants from each well were removed and 100 μl of BD Cytofix was added to each well. Cells were analyzed by flow cytometry, and the data collected were analyzed in FlowJo software. The percentage of fluorescent or bead+ cells and the median fluorescence intensity of the phagocytic cells were computed to determine the phagoscore.

### Quantification of plasma and tissue retinoid concentrations

Aliquots of previously isolated plasma were stored at -80°C, and retinoids were subsequently extracted as previously described by liquid-liquid extraction under yellow light (72–74). atRA was determined by liquid chromatography-tandem mass spectrometry with atmospheric pressure chemical ionization in positive-ion mode using an AB Sciex 5500 QTRAP hybrid triple-quadruple mass spectrometer (Foster City, CA) (72–74). ROL and RE were determined by HPLC/UV using an Aquity H-Class UPLC with a PDA detector (Waters, Milford, MA). Tissue retinoids were similarly quantified by first homogenizing frozen tissue in ground glass homogenizers (Kontes, size 22) in 0.5 to 1.0mL saline (0.9% NaCl) as has been previously described (72, 40). atRA was quantified with an API-4000 (Applied Biosystems). ROL and RE were quantified by HPLC/UV with an Alliance 2690 (Waters, Milford, MA). Calibration curves were determined for each analyte and utilized for retinoid concentrations. Plasma retinoids are expressed in moles per milliliter. Tissue retinoids are expressed as moles per gram of tissue.

### Stool sample collection, DNA extraction, and sequencing

Fecal samples were collected from all macaques at baseline (naïve), day 204 (day 21pi/week 3pi), and day 267 (day 84pi/week 12pi) and frozen at -80°C. Additionally, fecal samples from the vaccinated monkeys were also collected on Day 169/week 26 (2 weeks before challenge) to establish microbiome changes due to vaccination alone. Feces were later thawed, and DNA was isolated by spin column chromatography using a Stool DNA Isolation Kit (Norgen Biotek Corp, 27600) according to the manufacturer’s instructions. DNA was quantified using a GE SimpliNano spectrophotometer and aliquoted to be sent on dry ice to LC Sciences, LLC (Houston, TX, USA) for 16S ribosomal RNA (rRNA) sequencing. Universal primers were used to amplify V3+V4 for 16S rDNA fragment sequencing using the Illumina MiSeq platform. Output statistics were used to perform LEfSE to compare all groups.

### Antibody response

ELISA plates were first coated overnight with 1µg/ml (100uμl/well) HIV-clade C gp140 (strain 92BR025 #40251-V02H Sino Biologicals) and incubated. Similarly, to measure total IgG and IgA in CVL samples, separate plates were coated with 1.5µg/ml of polyclonal goat anti-monkey IgG + IgA + IgM (LS Bio) in coating buffer, 4°C overnight (100 µL/well). After incubation, the next day the plates were washed and blocked with (300 µL/well) blocking buffer at room temperature for 1hr. Plasma samples were diluted at 1:50, followed by 5-fold dilution steps, while CVL samples were diluted at 1:100, followed by serial 4-fold dilutions. Next, 100μl/well of plasma/CVL were added to the plates and incubated overnight after washing. Following this incubation, either goat anti-monkey IgA HRP (Rockland #617-103-006) or mouse anti-monkey IgG HRP (Southern Biotech #4700-05) 1:15,000 dilution was added. The plates were washed again, and then 100 µL/well of TMB substrate (A+B, KPL) was added, followed by a 30 min incubation. Finally, 100 µL/well stop solution was added, and the absorbance was read at 450nm.

### Statistical Analysis

Data were plotted using GraphPad Prism software and are represented as mean values. Two-way ANOVA was used for comparison of longitudinal viral loads between vaccinated and control animals. Mann-Whitney unpaired t-tests were used for specific time-point comparisons between vaccinated and control groups in all other analyses. In this study, p<0.05 has been considered statistically significant. For statistics, the anomalous monkey, A13R037, from the vaccinated group was excluded from viral load statistical analysis due to it being a suspected case of breakthrough infection.

## Data Availability Statement

The datasets generated and analyzed during this study are available within the manuscript.

## Acknowledgments

We thank the veterinarians and staff from the Comparative Medicine department at the University of Nebraska Medical Center for housing and assistance with animal procedures. HIV gp140 envelope glycoprotein (CH505TFchim.6R.SOSIP) (Duke University, credit Dr. Burton Haynes, Director, Duke Human Vaccine Institute) Figure 1, generated using BiRender.com. These studies were supported partially by R01AI129745; R01AI113883; R21MH113455; and R21AI114415 to SNB and philanthropic funds from Dr. Chokkavelu’s family to SNB research support.

## Author Contributions

SNB and VC designed the studies, obtained the required funding, and wrote and edited the manuscript. MT performed animal experiments and all virological assays and wrote a draft of the manuscript; KP and MJ helped in animal experiments; SJ performed all the microbiome studies and wrote/edited the manuscript; OA performed all immunological assays. JY and MK performed RA analysis in plasma and tissues. SA, KYH, and MAM performed antibody measurements; HG and DM performed neutralization assays; PW. JRB and FV provided regents and help in the design of the immunization DB, and GRK performed antibody assays. All authors read and edited the manuscript.

## Supplementary Figures

**Supplementary Figure 1.** Cell-associated viral DNA measured in different tissue compartments by ddPCR. (A) Reproductive tissues, (B) lymphatic tissues, and (C) and gut tissues. Bars indicate the median viral DNA copy number and individual values for vaccinated animals are represented in blue, while control animals are in red. The dashed line indicates the limit of detection for the assay (20 copies/mL).

**Supplementary Figure 2.** Gating strategy for CD4+ cells in PBMCs and various lymph node tissue cells. Briefly, dead cells were excluded based on zombie UV dye-positive expression. CD45+ cells were gated, followed by the generation of a total CD3+ T cell pool. From this, CD4+ T cells were gated out of the total cells and then further investigated to determine the extent of IFN-γ, CD107a, TNF-α, and IL-4 cytokine secretion following gag-specific stimulation compared to unstimulated (negative) controls. PMA and ionomycin were included as positive controls.

**Supplementary Figure 3.** Gating strategy for CD8+ cells in PMBCs and various lymph node tissue cells. Using zombie UV dye positive expression, dead cells were excluded. Then, CD45+ T cells were gated, and CD8+ cells were further obtained from the total CD3+ T cell pool. These cells were then evaluated to understand the extent of IFN-γ, CD107a, TNF-α, and IL-4 cytokine secretion following gag-specific stimulation compared to unstimulated (negative) controls. PMA and ionomycin served as positive controls.

**Supplementary Figure 4.** Mucosal vaccination is not associated with significant differences in all-trans retinoic acid precursors. Concentrations of retinol for (A) plasma and (B) duodenum and ileum tissue, and concentrations of retinyl esters for (C) plasma and (D) duodenum and ileum tissue.

**Supplementary Figure 5.** *Helicobacter* and viral dynamics. (A) The anomalous A13R037 experienced a rapid expansion of *Helicobacter* after vaccination, but before infection. (B) Within the vaccinated group, the relative abundance of *Helicobacter* after vaccination (Day 169) was predictive of acute plasma viral load at Day 204 (21dpi/week 3pi).

**Supplementary Figure 6**. Antibody responses in plasma and CVL fluids. We measured the gp140 specific as well as total IgA and IgG titers in control and vaccinated RMs. A-Plasma antibody responses; B-CVL antibody responses.

